# FLIMJ: an open-source ImageJ toolkit for fluorescence lifetime image data analysis

**DOI:** 10.1101/2020.08.17.253625

**Authors:** Dasong Gao, Paul R Barber, Jenu V Chacko, Md Abdul Kader Sagar, Curtis T Rueden, Aivar R Grislis, Mark C Hiner, Kevin W Eliceiri

**Affiliations:** Laboratory for Optical and Computational Instrumentation, Center for Quantitative Cell Imaging, University of Wisconsin, Madison, WI 53706, USA; UCL Cancer Institute, Paul O’Gorman Building, University College London, London, WC1E 6DD, UK; Department of Biomedical Engineering, University of Wisconsin, Madison, WI 53706, USA; Department of Medical Physics, University of Wisconsin, Madison, WI 53706, USA; Morgridge Institute for Research, University of Wisconsin, Madison, WI 53706, USA

**Keywords:** FLIM, Fluorescence Lifetime, Fluorescence imaging, image analysis, software, ImageJ

## Abstract

In the field of fluorescence microscopy, there is continued demand for dynamic technologies that can exploit the complete information from every pixel of an image. One imaging technique with proven ability for yielding additional information from fluorescence imaging is Fluorescence Lifetime Imaging Microscopy (FLIM). FLIM allows for the measurement of how long a fluorophore stays in an excited energy state and is affected by changes in its chemical microenvironment, such as proximity to other fluorophores, pH, and hydrophobic regions. This ability to provide information about the microenvironment has made FLIM a powerful tool for cellular imaging studies ranging from metabolic measurement to measuring distances between proteins. The increased use of FLIM has necessitated the development of computational tools for integrating FLIM analysis with image and data processing. To address this need, we have created FLIMJ, an ImageJ plugin, and toolkit that allows for easy use and development of extensible image analysis workflows with FLIM data. Built on the FLIMLib decay curve fitting library and the ImageJ Ops framework, FLIMJ offers FLIM fitting routines with seamless integration with other ImageJ components, and the ability to be extended to create complex FLIM analysis workflows. Building on ImageJ Ops also enables FLIMJ’s routines to be used with Jupyter notebooks and integrate naturally with science-friendly programming in, e.g., Python and Groovy. We show the extensibility of FLIMJ in two analysis scenarios: lifetime-based image segmentation and image colocalization. We also validate the fitting routines by comparing against industry FLIM analysis standards.

## Introduction

In the last thirty years, numerous advanced biological imaging techniques have allowed for the interrogation of biological phenomena at cellular and subcellular resolution. One of these powerful techniques is modern fluorescence microscopy, empowered by key inventions such as the laser scanning microscope and the use of fluorescent proteins. Among the range of modalities, fluorescence lifetime has been of particular interest in molecular imaging diagnostics due to its ability to probe the cellular microenvironment, sensitivity to changes in molecular conformations, and utility in interpreting phenomena such as Förster Resonance Energy Transfer (FRET) and physiological states including pH and hydrophobicity. Changes in the fluorescence lifetime can reveal important associations between the physical structure and the chemical microenvironment of a molecule. Based on this principle, Fluorescence Lifetime Imaging Microscopy (FLIM) has been adopted as a biological microscopy technique for investigating molecular level cellular behaviors. FLIM is now widely used in a variety of applications from measuring the metabolic state of differentiating stem cells (1) and intrinsic signatures of cancer cells (2) to measuring changes in lipid rafts (3) and FRET of signaling events in cell division(4). FLIM is available in two primary modes of operation, time-domain (5) and frequency domain (6), but also is compatible with a number of different microscopy configurations including wide-field (7), confocal (8), spinning disc (9) and multiphoton (10) microscopy. Several research groups have demonstrated super-resolution FLIM (11) and medical applications of FLIM, including ophthalmology and endoscopy (12). These emerging applications of FLIM continuously push further the innovation of FLIM technology including faster electronics and more sensitive detection. Despite these advances in biological applications and instrumentation, there has been a surprising lack in corresponding development in image informatics tools to directly support the FLIM imaging and analysis. Specifically, as a quantitative technique that generates image datasets, FLIM has the inherent need for powerful downstream image analysis software to interpret the results.

While more is needed, many of the recent advances in FLIM have largely been enabled by improved computation and software. Advanced software tools have not only allowed for FLIM electronics to be robustly controlled and capture short lifetimes, but to do so rapidly so that FLIM images in 3D (space) and 4D (space and time, or space and spectral) can be collected (13). Curve fitting algorithms have been developed that allow for robust fitting of two or more components. Several companies have made commercial packages for FLIM analysis, but these are closed source tools that are not transparent in their analyses and typically only support their own file formats. This makes sharing of approaches and FLIM data difficult while also limiting the usage of the features supported by the software. In recognition of the need for more transparent and customizable methods for FLIM analysis, there are new developments for new turnkey open analysis FLIM software tools such as FLIMfit from the Paul French’s group (14). However, most of these software packages are not designed with the rationale that FLIM results should be treated as images that can be segmented, statistically analyzed, or learned by upcoming newer machine learning algorithms. The separation of FLIM from other image analysis workflows has placed difficulty for biologists who would otherwise benefit from an easier image-based integration of FLIM.

To summarize, the scientific community would benefit from a more complete informatics approach addressing three specific and currently unmet needs:

1. an open and extensible FLIM algorithm library that supports most popular FLIM file formats and can be utilized and modified easily by a developer;
2. a turnkey FLIM analysis tool that uses the library and yet still can be used easily and directly by the bench biologist; and
3. the integration of FLIM analysis with versatile microscopy image analysis.

To address the first two needs, Barber et al. developed the Time-Resolved Imaging version 2 (TRI2), a freely available closed source FLIM analysis application equipped with a LabWindows graphical user interface (GUI) and basic image analysis capabilities (15). TRI2 was released to selected researchers in 2004, and from it, the core fitting algorithms were extracted to form the open-source FLIMLib library (see below).

The remaining need can be fulfilled by exploiting established open-source image analysis platforms. A great platform to integrate FLIM analysis is the open-source ImageJ (16) and its distribution for the life sciences, Fiji (17). ImageJ has long enjoyed use and adaptation by experimental biologists. However, FLIM workflows have largely been segregated from ImageJ due to lack of the necessary FLIM analysis functionality. Fortunately, the recent development of the ImageJ Ops framework (18) has laid a solid foundation for such a FLIM analysis toolbox. As a backbone of the next-generation ImageJ2 that powers Fiji, the ImageJ Ops framework supports the growth of ImageJ’s image processing and analysis power. In addition to offering hundreds of easy-to-use, general and efficient image processing operations such as segmentation algorithms, statistics, and colocalization methods across hundreds of file types, this framework provides programming interfaces for developers to extend the library. For example, the Ops-based plugins naturally cater to the needs of both bench biologists and advanced developers by allowing the former to easily use the Ops and the latter to build user-tailored applications for their own needs. Together with the seamless connections between Ops, the ImageJ Ops framework creates a suitable environment for developing modular, extensible image processing workflow components.

There were previous efforts by our group to integrate FLIM analysis into ImageJ known as the SLIM Curve (Spectral Lifetime Imaging Microscopy) plugin for ImageJ, which provided a graphical interface to FLIMLib. The SLIM Curve plugin was able to integrate FLIMLib’s full fitting functionalities with ImageJ while being accessible to bench biologists. However, the SLIM Curve plugin was built on legacy ImageJ 1.x data structures, which limited its extensibility and modularity compared to the ImageJ2 infrastructure.

We now have built on previous efforts in lifetime analysis (15,19) and spectral lifetime analysis (20) to build an ImageJ-centric toolkit, FLIMJ, that directly addresses these three needs. FLIMJ is an international collaboration between software developers and microscopists at the UCL Cancer Institute, London, UK (and formerly at the Gray Institute for Radiation Oncology and Biology at the University of Oxford, UK) and the UW-Madison Laboratory for Optical and Computational Instrumentation (LOCI) in the U.S. that leverages several existing software projects. When designing this FLIM analysis system, we recognized that current methods, such as those in MATLAB, may be difficult for the biological community to use. Further, many of these methods are also not attractive for developers because much of the published code, such as Numerical Recipes (21), may have restrictive licenses and a steep learning curve. We sought to develop a toolkit that would complement current commercial efforts and in fact directly support the image acquisition and image processing formats of these systems. We decided to focus our initial efforts on time-resolved FLIM data such as that collected from the PicoQuant and Becker & Hickl hardware systems, but the toolkit was designed to be flexible enough to support frequency or spectral domain in the future. Currently, spectral support is minimal, but this will be augmented as the toolkit evolves. This flexibility is supported by the use of the ImgLib2 (http://imglib2.net/) data container as widely adopted by almost all of the ImageJ plugins, which allows for the support of data of, in principle, unlimited dimensions and will handle FLIM data that includes additional channels of spectra and polarization.

As described below, the FLIMJ toolkit can either be invoked as an ImageJ Op or used from the graphical user interface (GUI) equipped plugin. One powerful advantage ImageJ can offer in FLIM analysis is segmentation. Looking at the lifetime distribution of a specific manually defined ROI can be often cumbersome or not even possible with commercially available FLIM analysis software packages. ImageJ not only has conventional manual segmentation but in addition, supports machine learning-based classification techniques (22), which can be exploited to use a training set to segment images in batch processing automatically. A number of other ImageJ features could be utilized with FLIM data including scripting, 3D visualization, and feature tracking, as well as a myriad of imaging file formats that can be supported through the use of the SCIFIO (http://scif.io/) infrastructures (23) to support a range of open and proprietary FLIM file formats. To our knowledge, this is the first FLIM analysis system that offers this range of flexibility and functionality.

The approach to use a core open-source software library brings the usual advantages of open-source that enable community contribution to and verification of the underlying code. It also enables the release of a ‘polished’ graphical-user-interface-driven plugin and allows use in other environments. In the following section a description of the toolkit is given, followed by how it can be extended for some more advanced uses.

## Methods

FLIMJ provides access to a variety of FLIM analysis techniques including standard nonlinear least-squares fitting in the form of the Levenberg-Marquardt (LM) algorithm and more advanced algorithms such as maximum likelihood, global, and Bayesian analysis that has been optimized for FLIM (15), as well as simpler methods such as the rapid lifetime determination (RLD) by integration (24), and frequency domain analysis via the method of phasors (25,26). In particular, the toolkit has the ability to account for an instrument response function (IRF, prompt, or excitation function) that distorts the pure exponential decay. Through iterative reconvolution, the LM algorithm extracts the true lifetime estimate from IRF distorted signals. The addition of new methods is also allowed and can inherit the standard library code interface as exemplified by the recently incorporated Bayesian algorithms (27). Similar to any other open or proprietary FLIM analysis software, FLIMJ consumes FLIM datasets of any accessible format (supported by Bio-Formats (28)) and outputs the results as images for further examination. By using ImageJ’s high-dimensional image data structure for both the input and output, FLIMJ ensures maximal compatibility with other parts of ImageJ, which can provide powerful preprocessing and post-analysis for the FLIM workflow (see Results).

The toolkit employs a modular design and comprises three major components: FLIMLib, FLIMJ Ops, and FLIMJ-UI. FLIMLib is the underlying library that contains an efficient C implementation of the algorithms. Based on the ImageJ Ops framework, FLIMJ Ops implements adapter ops for each of the algorithms in Java to dispatch the input data, invoke the corresponding C routines in FLIMLib, and collect the results. While FLIMJ Ops, together with the underlying FLIMLib deliver core functionality programmatically, FLIMJ-UI greatly improves the accessibility for bench biologists by providing an intuitive GUI based on the JavaFX framework to allow for the visualization and fine-tuning of the fit. In the rest of the section, we present a detailed description of each of the components.

### FLIMLib

FLIMLib is a cross-platform compatible library written in ANSI C with a Java API extension. With help from the current Maven-CMake building mechanism, the library can be compiled to run fast as a native executable on Linux, Windows, or macOS. As the weight-lifting component of FLIMJ, FLIMLib is equipped with a Java Native Interface (JNI) wrapper created by the SWIG framework (http://swig.org/), which offers efficient type conversion and data transfer between C and Java applications. However, more connectors can be added to make the library accessible to many high-level programmers using Python, MATLAB, C++, and C#, to name a few. Besides integrations with other languages, the library can also be compiled into a standalone command-line program with the intention that user interaction with the library could be in several forms according to the user’s choice. That is, the interaction could be through a graphical user interface such as with TRI2 or ImageJ, or could be through the command line in a scriptable form, or could be via a third-party framework such as MATLAB or R (http://www.r-project.org/). Full details on FLIMLib, including downloads, can be found on the project web site at https://flimlib.github.io/

The following fitting and analysis methods are currently present in the open-source library for lifetime data:

- ***Rapid Lifetime Determination (RLD)***: A method based on 3 integrals to determine a single average lifetime (24), including a variant that accounts for the IRF.
- ***Levenberg-Marquardt (LM) Non-Linear Least Squares Fitting***: A classical LM algorithm (29), the performance of which is modified by the noise model. Multi-exponential and stretched exponential analyses are built-in, others can be added, as are parameter fixing and restraining. There are variants with and without an IRF. Parameter error estimates are returned based on the fitting alpha matrix (21).

- ***Maximum Likelihood Estimation***: This variant of the LM algorithm is accessed by using a specific noise model. Several noise models can be chosen to influence the LM optimization via the chi-squared(χ^2^) parameter. The models are a) ‘***Constant’*** - every data point is assumed to have the same supplied variance. b) ‘***Given’*** - every data point can have an individual variance, given via a data array. c) ‘***Gaussian’*** - Variance for Gaussian distribution is used at all data points. d) ‘***Poisson’*** - Variance for Gaussian distribution is used with a lower limit of 15, this being the point where the Gaussian approximation begins to break down with Poissonian data. e) ‘***MLE’*** - Maximum likelihood estimation through the use of the Poisson equation (30).
- ***Global Analysis***: In some biological experiments, it is extremely advantageous to analyze data from a whole image or a series of images simultaneously to determine certain parameters with a high accuracy yet allow other parameters to remain variable at the pixel level to capture spatial variations. In this so-called ‘global analysis,’ some parameters can be globally determined while others remain local (31,32). This method is particularly useful when a biosensor based on FRET is in use (33), which usually exists in either an activated or deactivated state that is represented by different measured lifetimes by FLIM. Thus, these characteristic lifetimes can be determined globally, whilst the determination of the fraction of activated molecules can be determined locally (15). The library has built-in optimized functions for this type of analysis (19), that offer fast convergence and built-in optimization of initial parameter estimates. Methods for global analysis involving other generic functions (e.g., an exponential rise or non-exponential functions) are also included in the library.
- ***Phasor***: Transformation to phasor space for the calculation of a single average lifetime and graphical multi-exponential analysis (26).
- ***Bayesian Inference***: Lifetime estimation based on Bayesian inference offers higher precision and stability when faced with data with low photon counts (low signal-to-noise ratio). This algorithm acts by combining evidence from the photon arrival times to produce robust estimates of lifetimes and the potential errors in those estimates. In in vitro experiments, it was found that the precision was increased by a factor of two compared to LM fitting, or acquisition time could be reduced by a factor of two for an equivalent precision. The algorithms in the library can be used to estimate the IRF and exponential decay simultaneously or can be used to perform model selection between mono- and bi-exponential fitting models (27).

### FLIMJ Ops

FLIMJ Ops is a plugin built upon the ImageJ Ops framework (18,34) that connects FLIMLib and the ImageJ ecosystem. Based on the ImageJ Ops template, the plugin adapts the single-transient RLD, LMA, Global, Bayesian, and phasor analysis functionalities from FLIMLib into dataset-level fitting ops accessible from ImageJ. With help from the ImageJ Ops framework, FLIMJ Ops provides a concise yet flexible programmatic interface that can be easily included in a scripting workflow (see Results).

#### Ops API

Conforming to the organization convention of ImageJ Ops, the fitting ops are contained in the `flim` namespace. Specifically, by calling `flim.fit*` with `*` being `RLD`, `LMA`, `Global`, `Bayes` or `Phasor`, the user performs the corresponding analysis implemented by FLIMLib over the dataset on each individual pixel. While fully preserving FLIMLib’s granularity of control overfitting parameters such as noise model, initial estimation, and IRF, the Ops API provides support for dataset-level preprocessing operations, including pixel binning and ROI cropping. To optimize for the ease of use and prevent the need for a verbose argument list, some of the op arguments are marked as optional `(required=false)` at definition so that the programmer can ignore them in op calls. Upon invocation by name, the ImageJ Ops framework automatically inspects the number and types of arguments passed in, matches the appropriate fitting op, and completes the argument list by assigning sensible default values to optional parameters.

FLIM dataset analysis tasks greatly benefit from the parallelized fitting of individual pixels if they can be analyzed independently. This paradigm is employed by TRI2 and SPCImage by leveraging either the central processing unit (CPU) multithreading or graphics processing unit (GPU) acceleration. However, in terms of portability, CPU multithreading is considered more favorable. So far, all fitting ops in FLIMJ Ops, except for the global analysis op, by default run on multiple CPU threads in parallel. The parallelization relies on the ImageJ `ChunkerOp` utility for dispatching the workload.

### FLIMJ-UI

Oftentimes, it may be desirable for analytical software to integrate tools for visualizing results and to allow fine-tuning of the configurations. This is especially true for FLIM applications since the fitting results are usually sensitive to the settings, and manual setting of parameters such as initial values and decay interval range is required to obtain the optimal fit. The FLIMJ-UI is an ImageJ plugin created for such needs. Based on the SciJava Command framework, the plugin invokes FLIMJ Ops to carry out the computation and displays the fitted parameters alongside the decay curve with a JavaFX GUI. Like any of the ImageJ Ops, the plugin can be started through scripting, or the user may launch it from Fiji during an image analysis workflow.

### Notebooks for FLIM Analysis

The notebooks are a recent addition to the scientific processing scenario that helps one to bind the processing pipeline (codes) with the input/output aided with comments descriptively using markdown-style notes. Experiments and Image datasets can now be associated with processing routines pointing out dependencies and programming environments used for extracting results. Currently, notebooks are available to most scientific languages, including Mathematica, MATLAB, Python, R, Julia, Groovy, and others under tag names of Jupyter, BeakerX, and Zeglin notebooks. We supplement the FLIMJ Ops with a Groovy and Python notebook that can help a beginner to use fitting FLIM data in an interactive and data-analysis friendly way. Two example demo notebooks are provided with the FLIMJ Ops repository: 1) a groovy notebook running on the BeakerX kernel that accesses ImageJ ops directly, 2) a python Jupyter notebook that accesses ImageJ ops through the PyImageJ interface to invoke FLIMJ ops.

## Results

In this section, we demonstrate FLIMJ workflow and validate the results using separate software and simulated data. Two use-case scenarios are presented for image-segmentation and image-colocalization. a) Segmentation: FRET efficiency calculation of segmented tumorous tissue, b) colocalization of NAD(P)H, and antibody distribution for microglia. FLIMJ is shown to collaborate seamlessly and performantly with the central components of each workflow, as well as to have the potential to be integrated with more complex ones.

### Use Case I: Segmentation

This use case is based on the FLIMJ plugin for ImageJ, and relates to linking fluorescence lifetime processing to other advanced image processing plugins within ImageJ to accurately measure protein dimer information in cancer tissue. Many cancers are driven by the dimerization of the members of the HER/ErbB protein family (EGFR, HER2, HER3 etc.) at the cell surface. This discovery has led to the invention of targeted therapies that specifically disrupt the ability of these proteins to dimerize, and with some success; e.g., a monoclonal antibody for EGFR, cetuximab, is now used alongside chemotherapy for certain colorectal and head & neck tumors, and trastuzumab is used to target HER2 in some breast cancers. We have recently published the use of FLIM/FRET to detect HER family dimerization in archived patient tissue from breast and colorectal cancer studies and clinical trials (30,35) and correlate this with the response to targeted treatment.

One challenging aspect of these measurements in tissue is the need to segment the tumor tissue from surrounding stroma and other normal tissue components. This can be achieved through multi-exponential lifetime filtering (36), but it is often better to use independent information provided by the tissue morphology provided by imaging to segment the tumor. In ImageJ, we have created a pipeline that uses trainable Weka segmentation (22) on the intensity image in parallel to FLIMJ to provide tumor segmented lifetime statistics. These results from two serial sections, stained with the FRET donor and acceptor, and donor alone as a control, provide a FRET measure of protein dimerization. Additionally, training can also be done on lifetime images post-processing to help identify species. Figure 2 shows representative sections of breast cancer tissue from the METABRIC study (35) and calculated FRET scores of HER2-HER3 dimerization.

**Figure 1:**
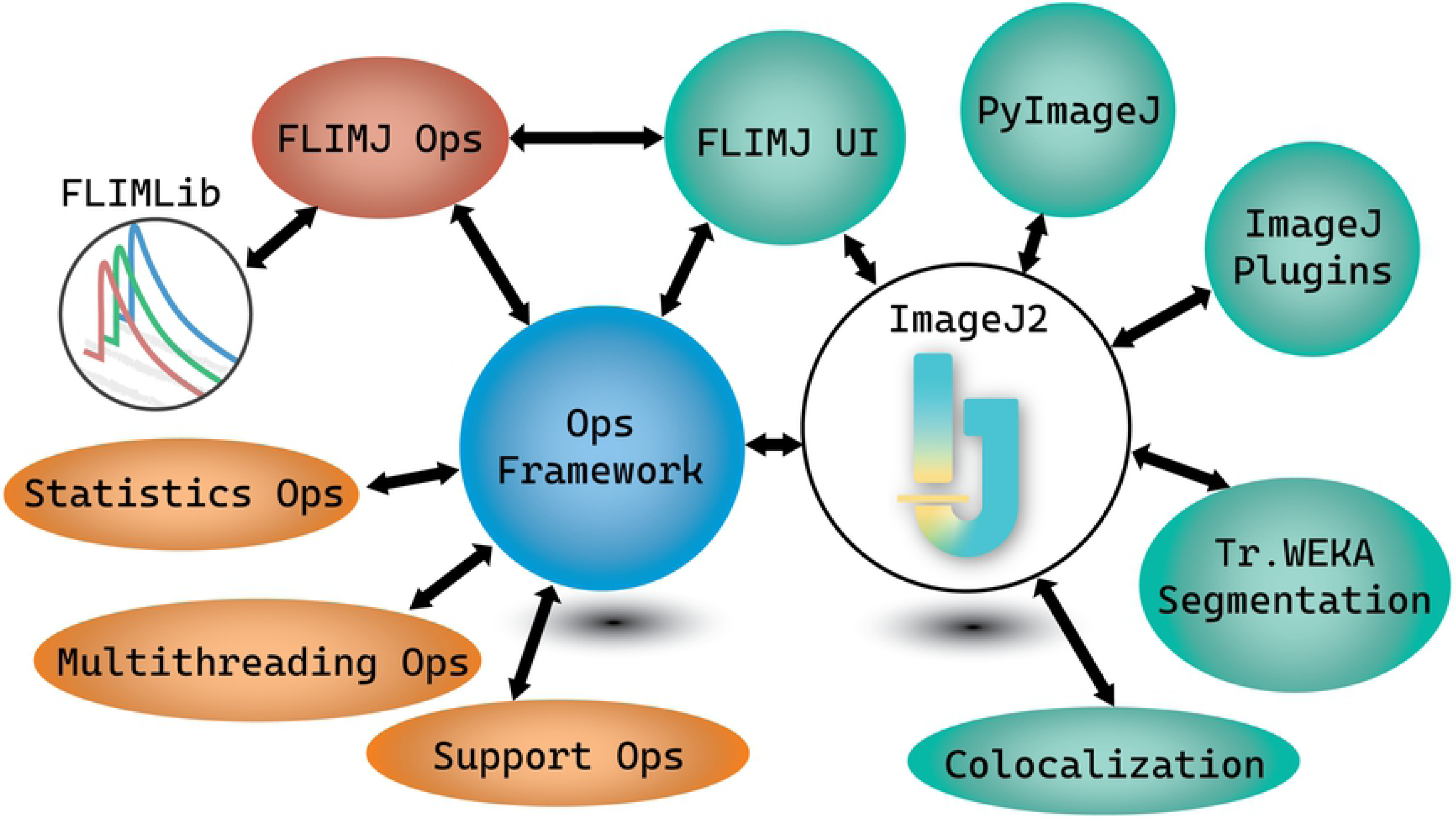
Relationships between components of FLIMJ and other parts of ImageJ2. FLIMJ Ops depends on FLIMLib and communicates with supporting ops through the Ops framework; other ImageJ plugins collaborate with FLIMJ Ops during FLIM workflows via FLIMJ-UI or directly through the Ops framework.

**Figure 2:**
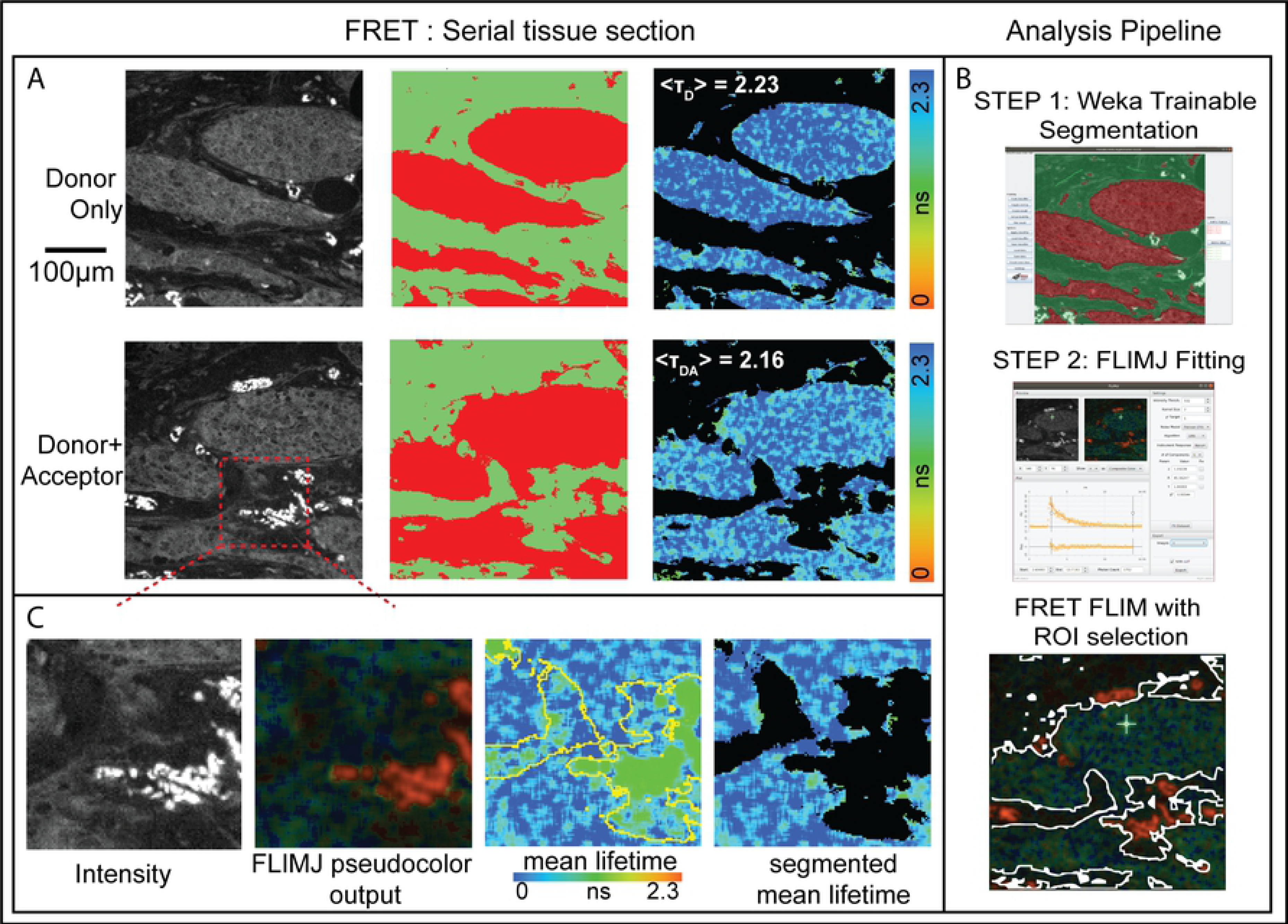
A) Breast cancer tissue from the METABRIC study was stained with antibodies: anti-HER3-IgG-Alexa546 (donor) and anti-HER2-IgG-Cy5 (acceptor). Serial sections were stained with donor + acceptor (FRET pair) and with donor alone (control). Average lifetime values can be determined for tumor from the two serial sections using FLIMJ after segmentation. The FRET Efficiency score is calculated for this region using the standard relationship.) B) Weka Trainable Segmentation plugin used to train the segmentation and showing a segmentation result (35). FLIMJ user interface showing a typical transient and fit from the tissue. We used LMA fitting with a mono-exponential model. C) Zoom into a smaller region. Composite image from FLIMJ showing lifetime information. Pure lifetime map with Weka segmentation shown in yellow. Segmentation result of the lifetime within the tumor with artifactual tissue removed. From T_D_= 2.23 ns and T_DA_= 2.16 ns, we estimate a FRET efficiency for the tumor area as 3.1%.

Samples were imaged on a customized “open” microscope automated FLIM system (36). Time-domain fluorescence lifetime images were acquired via time-correlated single-photon counting (TCSPC) at a resolution of 256 by 256 pixels, with 256-time bins and 100 frames accumulated over 300 seconds, via excitation and emission filters suitable for the detection of Alexa546 fluorescence (Excitation filter: Semrock FF01-540/15-25; Beam Splitter: Edmund 48NT-392 30R/70T; Emission filter: Semrock FF01-593/40-25). For technical convenience, those FLIM images were acquired through the emission channel of a UV filter cube (Long pass emission filter > 420 nm).

### Use Case II: Colocalization

This use case is based on the Fiji ROI colocalization plugin and links to fluorescence lifetime processing of autofluorescence images of microglia. Previously we and others have demonstrated the potential applicability of NAD(P)H FLIM in differentiating microglia functional state (37–39), and computational approaches to distinguish microglia cells (40). Based on our findings on the previous works, we expect that the hybrid method allowing lifetime estimation from raw decay data and subsequent colocalization analysis can be helpful in determining the effectiveness of FLIM based approaches in the identification of a specific cell type. The challenge lies in developing post-processing algorithms that yield maximum overlap between lifetime images generated from the endogenous signal and ground-truth images from the exogenous antibody fluorescence signal. To properly evaluate microglia identification with endogenous NADH signal(41), colocalization analysis can be of great benefit to quantitatively analyze overlap. Besides, there is concern that when GFP is used as a marker for cellular visualization and the NADH channel is used for lifetime analysis; there is a GFP signal bleed through to NADH channel, which can affect lifetime analysis(42). Colocalization analysis can help us identify pixels with higher GFP bleed through (43) and recalibrate the analysis. Fortunately, ImageJ includes the colocalization analysis plugin coloc-2, which is a great candidate for our use case that can be used in conjunction with lifetime analysis. This analysis approach can also be augmented by introducing another ImageJ-based analysis routine to perform post-processing on large scale datasets. Here, we explore the potential of FLIMLib fitting routines using FLIMJ connected with the ImageJ colocalization plugin (43). We use a ground truth generated from CX3CR1 GFP images and use the NADH FLIM signal from the same field-of-views. The NADH FLIM data is then fitted using two-component fit using FLIMJ, and the mean lifetime image is compared with ground truth image using colocalization plugin. The method is described in Figure 3. Panel C shows the overlaid image from mean lifetime (red) and antibody (green). Panel D shows the intensity histogram from the colocalization analysis using the coloc2 plugin and the colocalization was ~33%. This colocalization can help us find unwanted GFP signal bleed through to the NADH channel and help optimize the imaging configuration as well as find overlap between antibody channel and NADH when non-GFP antibody (iba1) is used.

**Figure 3:**
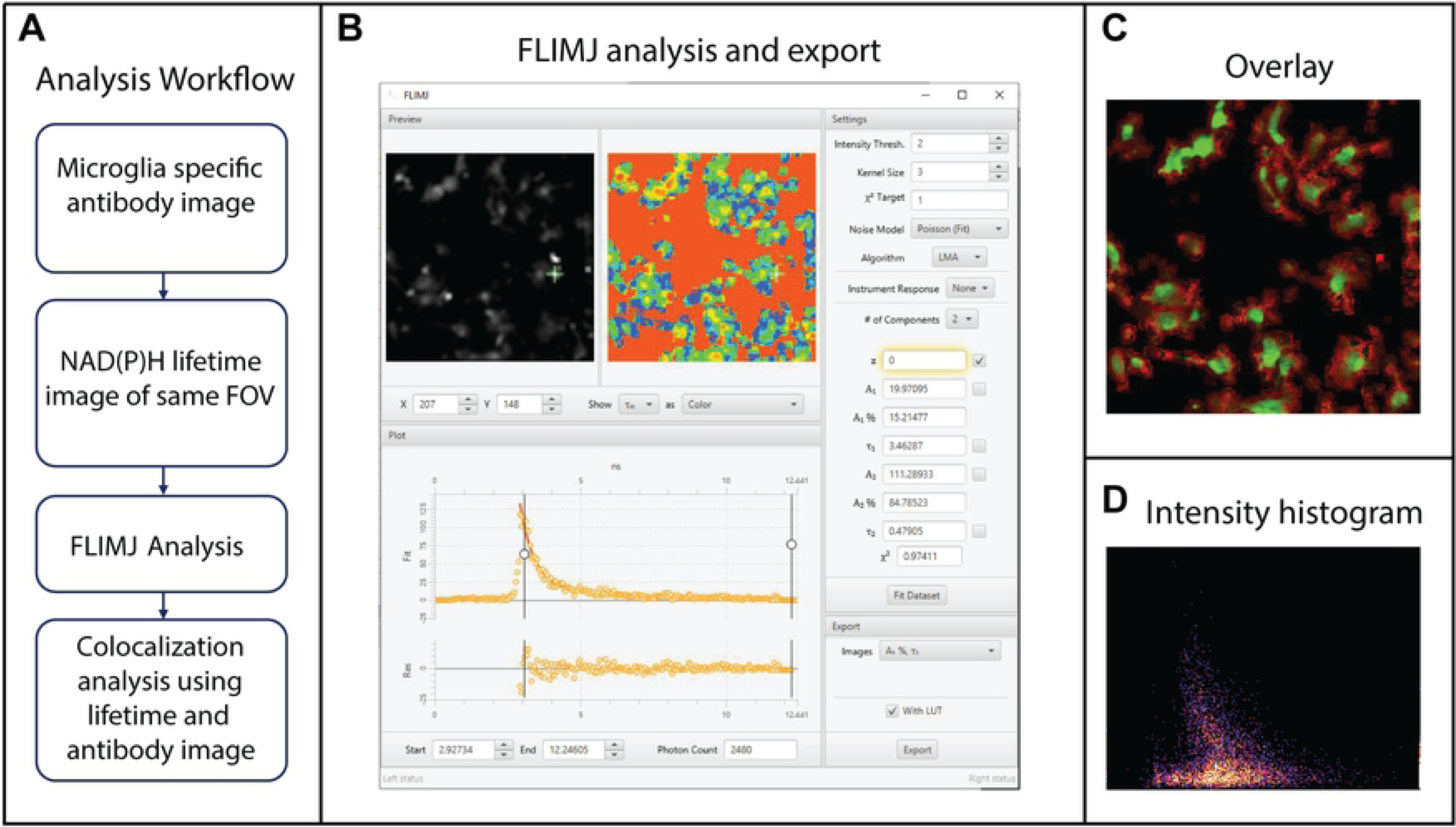
Microglia colocalization analysis using NADH FLIM. **A**) The analysis workflow describing the how microglia is visualized using specific antibody followed by NADH FLIM acquisition and FLIMJ analysis **B**) NADH FLIM data analysis using 2-component fit in FLIMJ-UI. **C**) Overlaid images of antibody (green) and lifetime image (red) **D**) Coloc2 analysis of mean lifetime and microglia antibody image

### Method Validation

We validated the lifetime value estimations of FLIMLib (see Figure: 4) using a FLIM image of a fluorescence standard: fluorescein in water with a known lifetime of 4.0 ns (44). The results were compared with the conventional fitting routine provided by the hardware vendor (SPCImage) and using the FLIMJ/FLIMLib routine. The data were analyzed using a 3×3 kernel and fit to a single component model in LMA setting for both analyses. The vendor’s routine results in an identical distribution to FLIMLib fitting results. This fitting results shown here are derived by setting the offset value to fixed numbers and fitting for the two parameters: lifetime and amplitude. The average fitting time for SPCImage and FLIMJ is identical for similar fitting parameters. There is an apparent change in speed when spatial binning is changed. FLIMJ performs spatial binning by convolving the input data with the kernel using FFT, and this operation is separate from the fitting, while SPCImage calculates the kernel every time within the fitting routine. The calculation times were computed for `SPCImage 7.4` and `flimj-ops 2.0.0` on the same computer. Neither of these comparisons used GPU optimization for testing, which could be advantageous for fitting large image datasets with homogenous fitting parameters.

We tested different fitting routines available within the flimj-ops framework (Refer to Figure 5). Two data sets were simulated for testing: 1) one component model and 2) a two-component model. The one component model was used to compare results from LMA, Phasor, and RLD methods. Fit results were plotted against the ground truth values, and a linear ~1:1 relation was seen between these three commonly used fitting methods. We also tested the phasor plots and found the phasors for single component lifetime curves fit on the universal circle. The universal phasor circle is based on the fitting range. In the simulations, we used a 117 MHz universal circle to match the range of fitting instead of the more common 80 MHz plot (matched to the repetition rate of the laser used)(45). Two significant advantages of our library, when compared to other available commercial software bundles, is its ability to use the Bayesian lifetime estimation method and Global fitting routines (31,46). We validated the `fitBayes` function on the same dataset. However, Bayesian estimates work best at low photon images such as auto-fluorescence-based FLIM and similar low-light FLIM experiments.

**Figure 4:**
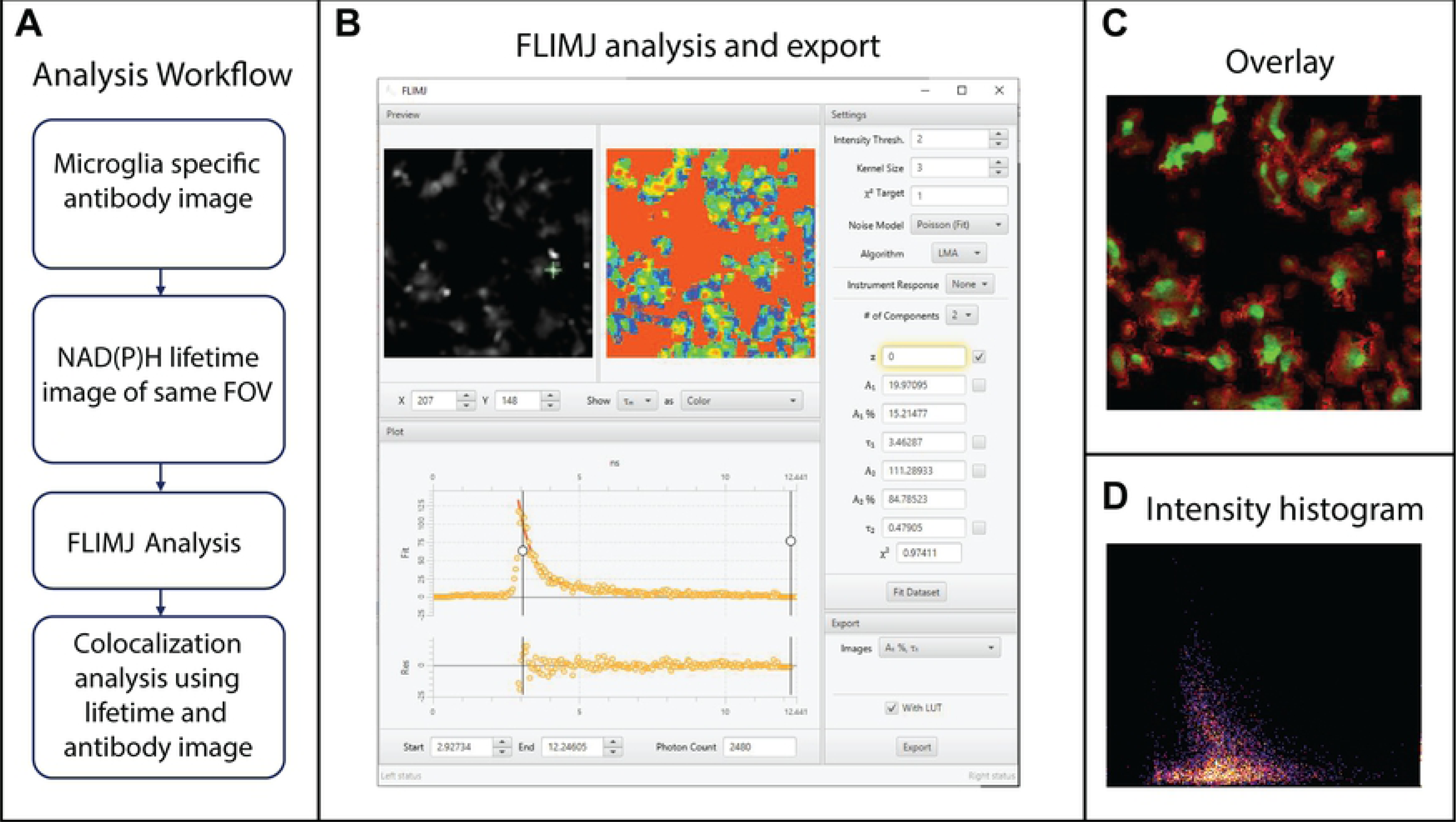
Validation of FLIMLib LMA lifetime estimation against hardware vendor-provided software with a solution of fluorescein in water. The phasor plot for the data is also shown as a proof of principle of the FLIMLib Phasor function for fit-less estimation of lifetime parameters. The two parallels of lifetime histograms and phasor plots are the current laboratory standards for FLIM analysis. The phasor is plotted on a universal circle derived from the endpoints of the fit-range (117 MHz).

**Figure 5:**
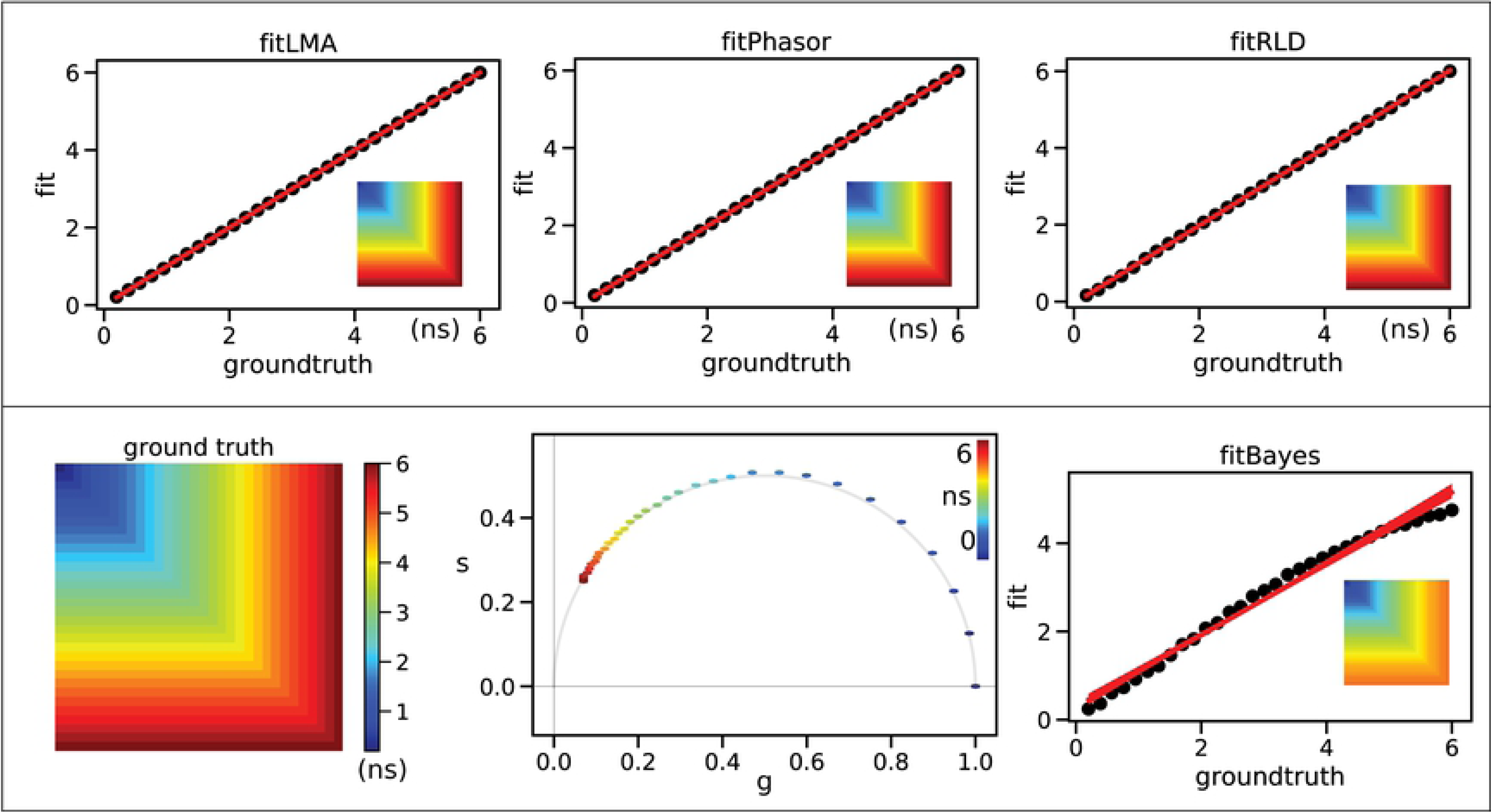
FLIM Validation using a simulated dataset for 1-component fitting. The lifetime values range from 0.2ns to 6.0 ns that fit within the fitting range of 10ns. The top panel shows a comparison of LMA, Phasor, RLD fitting; and the bottom panel shows the ground truth, phasor plot, and Bayesian results. The inset panels show the estimated lifetime maps.

For validating Global fitting, we simulated a spatially varying two-component model Refer Fig. 6). We demonstrate the fitting results from a data simulated by two fluorescent species (A and B) with fixed lifetime values (0.4ns and 2.1ns), with spatially varying fractions. These values were chosen as an example of widely used autofluorescence FLIM analyses of NAD(P)H and FAD (47,48). In the FLIM data shown, top-left is 100% species A and bottom right is purely species B. The image is made of 128×128 pixels. We tested all the available fitting models on this dataset to estimate the time taken by each method (without any spatial binning). The measured timings were: `fitLMA` required ~1 second, `fitGlobal` ~0.5 seconds, `fitRLD` ~0.16 seconds, `fitPhasor` ~0.27 seconds, and `fitBayes` ~0.95 sec *(using 1-component. fitBayes is currently implemented only for 1-comp analysis)*.

**Figure 6:**
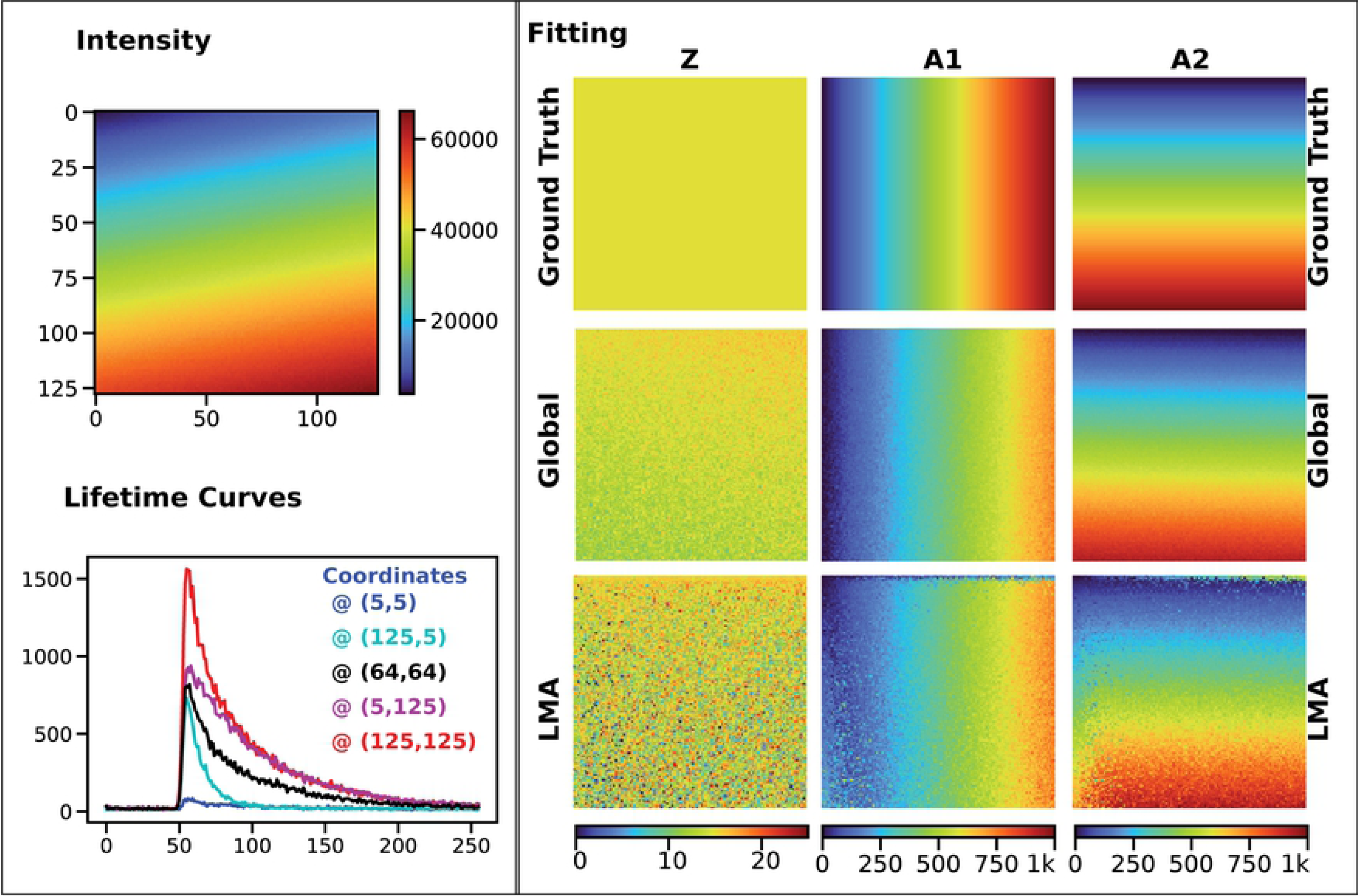
FLIM Validation using a simulated 2-component lifetime data. The left panel shows the simulated data with fixed lifetime values 2.1ns and 0.4ns. Five sample lifetime curves are shown to demonstrate the variation in intensity levels. The right panel compares the Global fitting routine and LMA. The color scales are the same between panels of each parameter. Both LMA and Global fitting does reproduce the fraction of two species, but we find global fitting yields lesser noise and works twice as fast. (This dataset is provided in the <u>SCIFIO</u> sample datasets or https://samples.scif.io/Gray-FLIM-datasets.zip.)

## Discussion

In this paper, we presented FLIMJ, an open-source toolkit for curve fitting and analysis of lifetime responses. We demonstrated how it could be integrated with a variety of ImageJ workflows, including segmentation, colocalization, and cross-language analysis for Python. The use of the toolkit is also possible from other languages such as JavaScript, Groovy, or the R statistics package.

The FLIMLib library includes a range of fitting routines for lifetime data based on Levenberg-Marquardt and Maximum Likelihood methods as well as analysis tools such as the phasor method and rapid lifetime determination from the area under the curve. With support from the ImageJ Ops framework, FLIMJ Ops is able to extend the usage of FLIMLib functions beyond plain curve fitting and seamlessly incorporates them with image analysis workflows.

The described analysis tools enable a wide range of FLIM experiments. Such as the simple detection of a change in lifetime due to a change in the chemical environment (49), or viscosity changes (32). As well as more advanced experiments such as the detection of FRET. Biologically useful extensions are also possible based on these core algorithms, such as global analysis, to increase signal to noise ratio in lifetime invariant systems, support plane analysis to determine ‘confidence’ in fitted parameters, and model selection to help selection of the most appropriate model for the data.

Although FLIMJ has been successful at integrating FLIM analysis into ImageJ as an image analysis workflow, several improvements can be made. This may be mitigated by introducing global optimization algorithms before and/or after the fit. Further, although the fitting routines have been verified with data from our laboratory, more validation can be done as users start to use FLIMJ on theirs collected from different systems. We also plan to improve FLIMJ Ops and FLIMJ-UI in terms of usability. Until the current stage of development, more of the effort has been put on implementing new features than on demonstrating the existing ones. While the FLIMJ Ops landscape is being finalized, we will shift our focus towards updating and creating demonstrative tutorials using the ImageJ Wiki page or Jupyter Notebooks. Other usability improvements may include allowing saving of fitting configurations as a workspace file as in TRI2, implementing batch-fitting in the GUI, and packaging FLIMJ as standalone runnable for those without access to ImageJ. Much of the future work will also focus on further extending the functionality of FLIMJ.

Development and distribution of FLIMLib, in particular, will be aimed at the simultaneous analysis of spectral, lifetime, and polarization information that is now routinely captured in some laboratories (20,50,51). This is an area in which novel advanced analysis is greatly needed, especially as photon numbers from biological samples are limited (52) and the addition of more dimensions of measurement (i.e. time, spectral, polarization) can result in a very small number of photon counts per measurement channel. Although the current implementation of FLIMLib has been able to deliver a decent throughput, it can still be further optimized for speed and simplicity by depending on open-source libraries such as GSL and Boost, which yields benefit to throughput-demanding applications including machine learning-assisted FLIM analysis. Lastly, as Fiji continues to be optimized for speed and performance, and explores parallelization and possible GPU based applications, these are areas where improved FLIM analysis performance can be evaluated as well.

## Data Availability

All relevant data used in this manuscript are provided in the SCIFIO sample datasets repository accessible at https://scif.io/images. The specific FLIM data can be accessed directly at https://samples.scif.io/Gray-FLIM-datasets.zip. The GitHub repository for the library and example notebooks are available at https://flimlib.github.io.

## Acknowledgements

We acknowledge B. Vojnovic, S. Ameer-Beg and J. Gilbey for their significant contributions to the development of the original algorithms in TRI2, J. Nedbal for the MATLAB wrapper to FLIMlib and M. Rowley and T. Coolen for the development of the Bayesian fitting algorithms. We also thank T. Gregg, M. Merrins and B DeZonia for their useful input on the original version of the program.

